# Transcriptomic classification of pituitary neuroendocrine tumors causing acromegaly

**DOI:** 10.1101/2022.07.26.501638

**Authors:** Julia Rymuza, Paulina Kober, Natalia Rusetska, Beata J. Mossakowska, Maria Maksymowicz, Aleksandra Nyc, Szymon Baluszek, Grzegorz Zieliński, Jacek Kunicki, Mateusz Bujko

## Abstract

Acromegaly results from growth hormone hypersecretion caused by somatotroph pituitary neuroendocrine tumor (PitNET). Our molecular profiling revealed that acromegaly-causing tumors form three distinct transcriptomic subgroups with different histological/clinical features. Transcriptomic subtypes of somatotroph tumors differ in the expression levels of numerous genes including those involved in hormone secretion and genes with known prognostic value. They can be distinguished by determining the expression of marker genes. Transcriptomic group 1 includes ∼20% of acromegaly patients with *GNAS* mutations-negative, mainly densely granulated tumors with *NR5A1* (SF-1) and *GIPR* co-expression. Group 2 tumors are the most common (46%) and include mainly *GNAS*-mutated, densely granulated somatotroph and mixed PitNETs. They have significantly smaller size and express favorable prognosis-related genes. Group 3 includes predominantly sparsely granulated somatotroph PitNETs with low *GNAS* mutations frequency causing ∼35% of acromegaly cases. Ghrelin signaling is implied in their pathogenic mechanism, they have unfavorable gene expression profile, and invasive growth rate. Since a subgroup of somatotroph tumors have high *NR5A1* expression, using SF-1 as classification marker specific to gonadotroph PitNETs could be reconsidered.

## 1 Introduction

Acromegaly is a severe and life-threatening disease caused by persistent excess of the growth hormone (GH) which stimulates synthesis and secretion of the insulin-like growth factor-1 (IGF-1). In the majority (95%) of patients acromegaly is caused by sporadic GH-secreting pituitary neuroendocrine tumor (PitNET). High IGF-1 level promotes cell proliferation, inhibits apoptosis, and causes most of the clinical symptoms of acromegaly, ranging from subtle to severe - limb hypertrophy, soft tissue edema, arthralgia, prognathism and hyperhidrosis to frontal bone hypertrophy, diabetes mellitus, hypertension, respiratory and heart failure. Excessive body growth and gigantism may develop when somatotroph tumor develops in young patients before closing the epiphyses of long bones (Melmed, 2006).

Acromegaly is a particular disease entity that is generally treated with somatostatin analogs and surgery. However, PitNETs that cause clinical symptoms are quite heterogenous, varying in terms of pathomorphological characteristics, radiological imaging results, invasiveness, and molecular features (Asa *et al*, 2017; Akirov *et al*, 2019). Diversity of these tumors was already addressed in the current histopathological WHO classification of PitNETs (Asa *et al*, 2022a). Most of somatotroph tumors expresses GH and PIT-1 transcription factor and are further divided into sparsely and densely granulated (SG and DG, respectively) with electron microscopy-based evaluation or anti-cytokeratin staining. These two types of somatotroph tumors differ in MRI features and clinical course of the disease (Swanson *et al*, 2021). Acromegaly can also be caused by mammosomatotroph and mature plurihormonal tumors, characterized by expression of GH and additional hormones (prolactin (PRL) or PRL and thyroid-stimulating hormone (TSH), respectively). Addidtionally it is caused by mixed somatotroph-lactotroph tumors (composed of two distinct populations of somatotroph and lactotroph tumor cells), immature PIT-1-lineage tumors and acidophil stem cell tumors (Asa *et al*, 2022a). SG somatotroph PitNETs and plurihormonal PIT-1 positive PitNETs are categorized as high-risk tumors due to characteristics of aggressiveness including invasive growth and higher recurrence rate (Kontogeorgos *et al*, 2022).

The most common molecular changes in somatotroph tumors are activating mutations in *GNAS* gene, whichencodes stimulatory subunit of heterotrimeric G protein. These mutations are present in approximately 40% of somatotroph PitNETs (Efstathiadou *et al*, 2015). They cause hyperactivation of cAMP-dependent pathways and, consequently, both increased secretion of GH and proliferation of somatotropic cells (Lania & Spada, 2009). Clinical significance of *GNAS* mutations is unclear, however, certainly, they are more common in DG than in SG tumors (Larkin *et al*, 2013).

Recently, a subset of somatotroph PitNETs with elevated expression of gastric inhibitory polypeptide receptor (GIPR) was identified (Regazzo *et al*, 2017; Occhi *et al*, 2011). In these tumors *GIPR* expression is considered to additionally stimulate cAMP pathway causing paradoxical GH response to glucose intake (Hage *et al*, 2021). These tumors were suggested to constitute a separate molecular subgroup as they differ in epigenetic profile and *GIPR* high expression is mutually exclusive with *GNAS* mutations (Hage *et al*, 2019).

Considering the complex nature of clinical and pathological spectrum of PitNETs causing acromegaly we aimed to investigate genes expression in somatotroph tumors to verify whether transcriptomic profiles correspond to current histological/clinical classification.

## 2 Materials and Methods

### Patients and tissue samples

This study included 134 patients with biochemically confirmed acromegaly that were treated with transsphenoidal surgery in two specialized centers: Department of Neurosurgery, Military Institute of Medicine, Warsaw and Department of Neurosurgery, Maria Sklodowska-Curie National Research Institute of Oncology, Warsaw in years 2013 – 2020. Diagnostic critera of acromegaly based on clinical characteristics, increased serum IGF-1 levels and non-suppressible GH after Oral Glucose Tolerance Test (OGTT), in patients with no OGTT contraindications were used. All patients received somatostatin receptors ligands (SRLs) treatment (octreotide or lanreotide) before surgery following the recommendations of Polish Society of Endocrinology (Bolanowski *et al*, 2019). Invasive growth was determined based on preoperative MRI using Knosp classification. Tumors scored with Knosp grades 0-2 were considered noninvasive while those with Knosp grade 3 and 4 were considered invasive (Knosp *et al*, 1993).

Each tumor sample was divided in three parts. One of them was snap frozen in liquid nitrogen and stored for molecular analysis and the other two were preserved for histopathological evaluation, including immunohistochemical staining and ultrastructural analysis with electron microscopy. Pathomorphological diagnosis was based on evaluation of the immunoexpression of pituitary hormones (GH, PRL, ACTH, TSH, FSH, LH, α-subunit) and Ki-67, as well as assessment of ultrastructural status (sparsely *vs*. densely granulated tumors). Expression of PIT-1 transcription factor was confirmed with immunohistochemical staining retrospectively since large proportion of the tumors was originally diagnosed with WHO 2014 criteria that did not comprise evaluation of transcription factors. Overall patients’ characteristics are presented in Table 1.

**Table 1.**
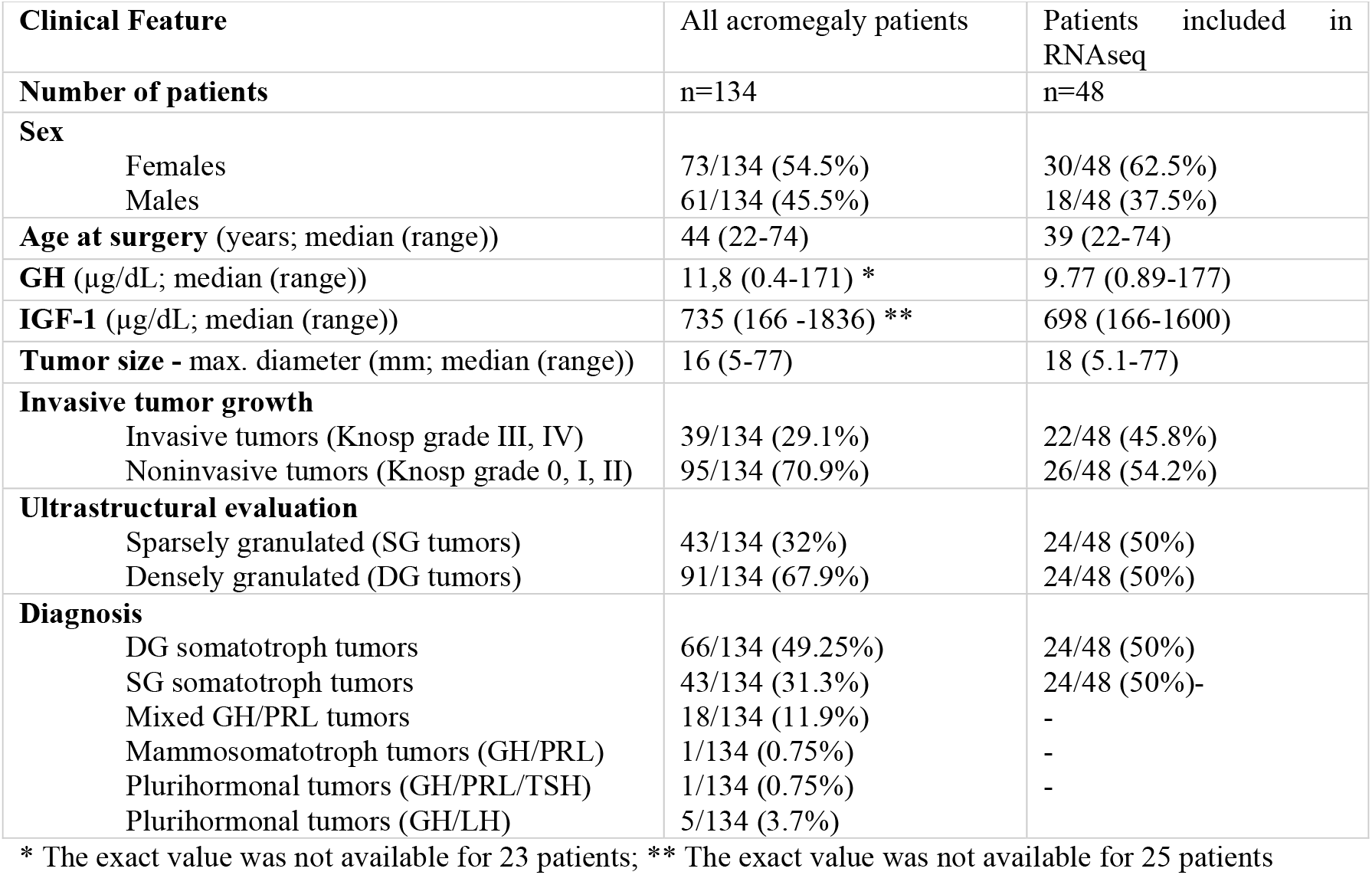
Summary of demographical and clinical features of patients with acromegaly

The study was approved by the local Ethics Committee of Maria Sklodowska-Curie National Research Institute of Oncology in Warsaw, Poland. Each patient provided informed consent for the use of tissue samples for scientific purposes.

DNA and total RNA from tumor samples were isolated with AllPrep DNA/RNA/miRNA Universal Kit (QIAGEN) and stored at -70°C.

### Testing for GNAS mutation status

The presence of *GNAS* point mutation (exons 7 and 8) was assessed with Sanger sequencing in 134 tumor samples. DNA was PCR amplified with FastStart Taq DNA Polymerase (Roche Diagnostics, Mannheim, Germany) using GeneAmp 9700 PCR system (Applied Biosystems, Foster City, CA, USA). PCR product was purified with ExoStar (GE Healthcare Life Sciences, Pittsburgh, PA, USA), labeled with BigDye Terminator v.3.1 (Applied Biosystems) and analyzed by capillary electrophoresis using ABI PRISM 3300 Genetic Analyzer (Applied Biosystems). PCR primers’ sequences are provided in Supplementary Table 1.

### RNA sequencing

Forty-eight tumor somatotroph PitNET samples including 40 *GNAS* wild type (*GNAS*wt) and 8 *GNAS* mutated (*GNAS*mut) were subjected to RNA sequencing. Library preparation was performed with 1 μg RNA from each sample using NEBNext Ultra II Directional RNA Library Prep Kit for Illumina. NEBNext rRNA Depletion Kit was used for ribosomal depletion. The quality of libraries was assessed using the Agilent Bioanalyzer 2100 system (Agilent Technologies, CA, USA). Libraries were then sequenced on an Illumina NovaSeq 6000 platform, and 150-bp paired-end reads were generated. Aminimum of 30 M read pairs per sample were generated. Sequencing was performed by Eurofins Genomics service.

### Analysis of RNAseq results

Quality control of raw reads was conducted using FastQC (Andrews *et al*, 2015). Raw reads were mapped to the human reference genome GRCh37/hg19 ref with HISAT2 (Kim *et al*, 2015). The raw unnormalized count matrix was generated using featureCounts (Liao *et al*, 2014) with gene features from GENCODE (v39) and imported to DESeq2 (Love *et al*, 2014). Low-expression genes (genes with less than five sequencing reads in less than 25% of samples) were excluded from further analysis. Filtered matrix was normalized using DESeq2 (Love *et al*, 2014) and used for sample clustering with k-means algorithm (R package cluster (Maechler *et al*, 2018)) and hierarchical clustering (Manhattan distance and ward.D agglomeration, R library stats). Analysis of genes differentially expressed between clusters (transcriptomic groups) was performed using DESeq2 (Love *et al*, 2014). Differentially expressed genes (DEGs) were defined as those with adjusted p-value < 0.05 and |FC|>2. Gene set enrichment analysis (GSEA) was conducted with fgsea (Sergushichev, 2016). Additionally, marker genes for each group were detected using R package MGFR.

### qRT-PCR gene expression analysis

One microgram of RNA was subjected to reverse transcription with Transcriptor First Strand cDNA Synthesis Kit (Roche Diagnostics). qRT-PCR reaction was carried out in 384-well format using 7900HT Fast Real-Time PCR System (Applied Biosystems) and Power SYBR Green PCR Master Mix (Thermo Fisher Scientific) in a volume of 5 μL, containing 2.25 pmol of each primer. The samples were amplified in triplicates. *GAPDH* and *SDHA* served as reference genes. Delta Ct method was used to calculate relative expression level with geometric mean of reference genes Ct value for normalization. PCR primers’ sequences are presented in Supplementary Table 1.

### Statistical analysis and data visualization

Two-sided Mann–Whitney U-test was used for analysis of continuous variables. The Spearman correlation method was used for correlation analysis. Significance threshold of α = 0.05 was adopted. Data was analyzed and visualized using GraphPad Prism 6.07 (GraphPad Software) and R environment. R libraries such a as ggplot2 (Gómez-Rubio, 2017) and plotly (Li & Bilal, 2021) were used for visualization. Moreover, scaled normalized RNAseq read counts were visualized on KEGG pathway graph using pathview (Luo & Brouwer, 2013).

## 3. Results

### 3.1 Incidence of GNAS mutations

First, we determined *GNAS* mutational status in tumor samples in the entire cohort of patients. The missense mutations were identified in 52/134 (38.8%) patients. Forty patients harbored mutations in exon 8 including 35 variants c.C601T:p.R201C, 4 c.A680G:p.Q227R and 2 c.C601A:p.R201S. Twelve patients had mutations in exon 9 including 7 variants c.A680AT:p.Q227L and 5 c.A680G:p.Q227R. We did not observe significant relationship between *GNAS* mutation and demographical/clinical features including patients’ age, gender, pathological diagnosis, invasiveness status and tumor size.

### 3.2 Whole transcriptome analysis

Forty-eight tumor samples (8 *GNAS*mut and 40 *GNAS*wt tumors) were successfully processed. An average 87,347,293 reads per sample were generated with an average 90.79% reads mapped to UCSC hg19 reference genome. The sequencing reads were mapped to 19,631 human protein-coding genes, and 16,096 mapped protein-coding genes remained for inclusion in subsequent analyses after low-expression genes were filtered out.

Data-dimensionality reduction analysis including principal component analysis (PCA) and uniform manifold approximation and projection (UMAP) clearly indicated the presence of three separate transcriptomic groups of tumor samples with groups 1 and 2 being more similar to each other than to the third group (Figure 1A). The same pattern was observed in hierarchical clustering analysis where three basic branches of clustering tree corresponded to transcriptomic groups observed in PCA and UMAP results (Figure 1B). Nearly the same clustering results were observed regardless of the number of differentially expressed genes included in the analysis. The results for 1%, 10%, 20% of most differentially expressed genes are presented in Supplementary Figure 1. The preliminary analysis of clinical data for discovery set of 48 samples revealed notable differences between three identified transcriptomic groups. Group 1 included *GNAS*wt DG tumors (except for one *GNAS*wt, SG tumor), group 2 included mainly DG tumors with a high proportion of samples with *GNAS* mutation, while group 3 included basically DG tumors with a low percentage of *GNAS*mut ones.

**Figure 1.**
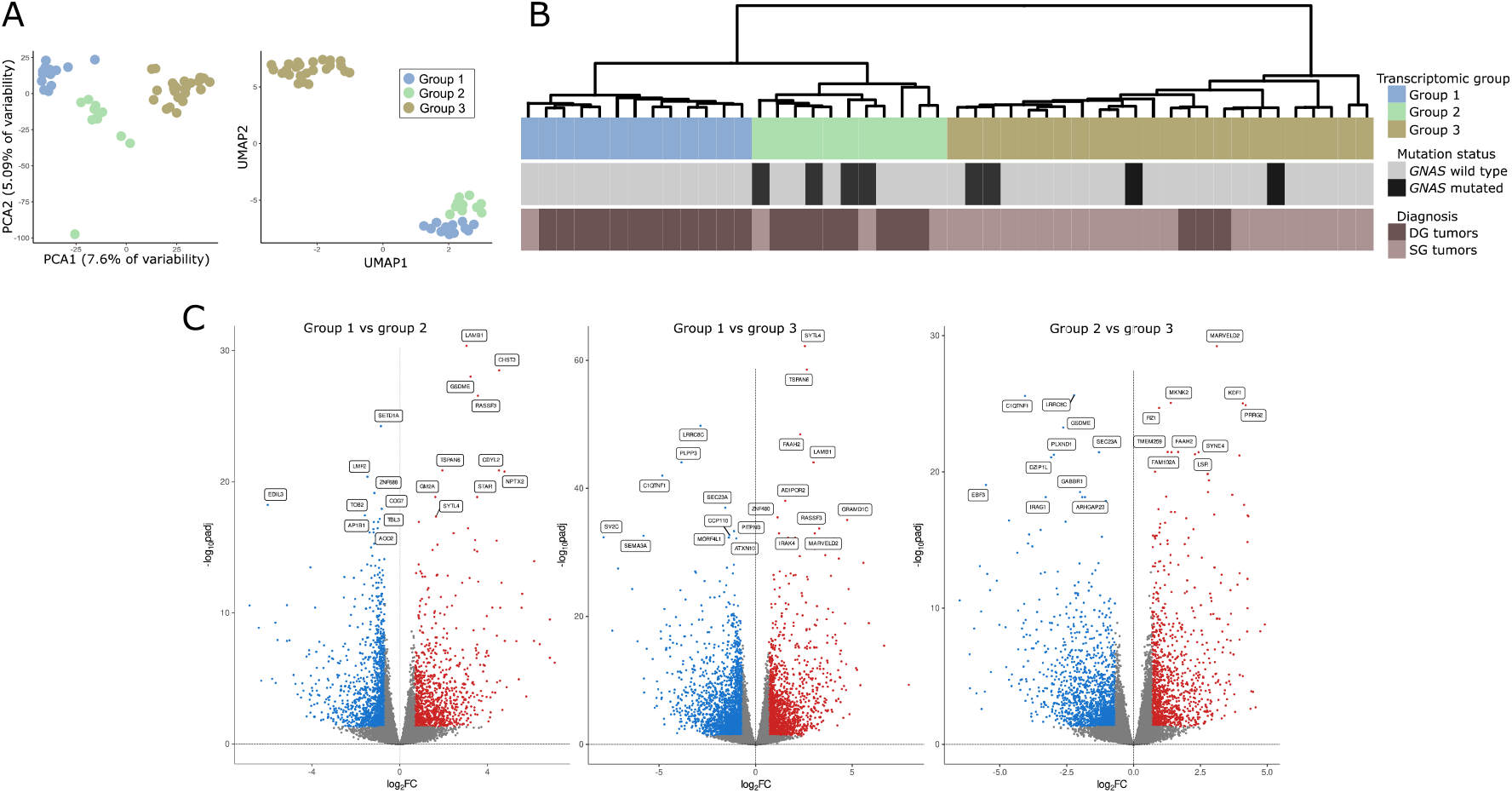
Gene expression in somatotroph tumors. A. Principal component analysis (PCA) and uniform manifold approximation and projection (UMAP) results based on the expression data for the entire set of genes that indicate the presence three transcriptomic groups of somatotroph tumors. B. Hierarchical clustering of somatotroph PitNETs according to the expression data for the entire set of genes, presented with basic diagnostic data and *GNAS* mutation status, DG stands for densely granulated somatotroph PitNET, SG stands for sparsely granulated somatotroph PitNET. C. Comparison of genes expression in pairs of particular transcriptomic groups.

We determined the genes that are differentially expressed between each of the transcriptomic groups by comparing each group with the remaining groups separately (group 1 vs group 2, group 2 vs group 3 and group 2 vs group 3). We found that paired groups differ in the expression of high number of genes. Specifically, 1,007 differentially expressed genes (DEGs) that met criteria |FC|>2 and adjusted p-value < 0.05 were found when comparing groups 1 and 2; 2,403 DEGs were identified when comparing groups 1 and 3 while there were 1,685 DEGs in comparison of groups 2 and 3. Results of differential analysis are presented in Figure 1B. The lists of differentially expressed genes are reported in Supplementary Table 2.

The functional implications of the difference in gene expression were investigated with GSE analysis with Gene Ontology (GO) including GO Biological Processes and GO Molecular function. The analysis resulted in the identification of a large number of significantly enriched GO terms. The most significantly enriched GO Biological Processes, according to the highest significance level were the terms related to G-protein signaling, ion transport, cellular adhesion and differentiation while the most enriched GO Molecular function terms were those related to signaling and ion transport. The top 10 most enriched terms for each comparison are presented in Figure 2, all significantly enriched terms are listed in Supplementary Table 3.

**Figure 2.**
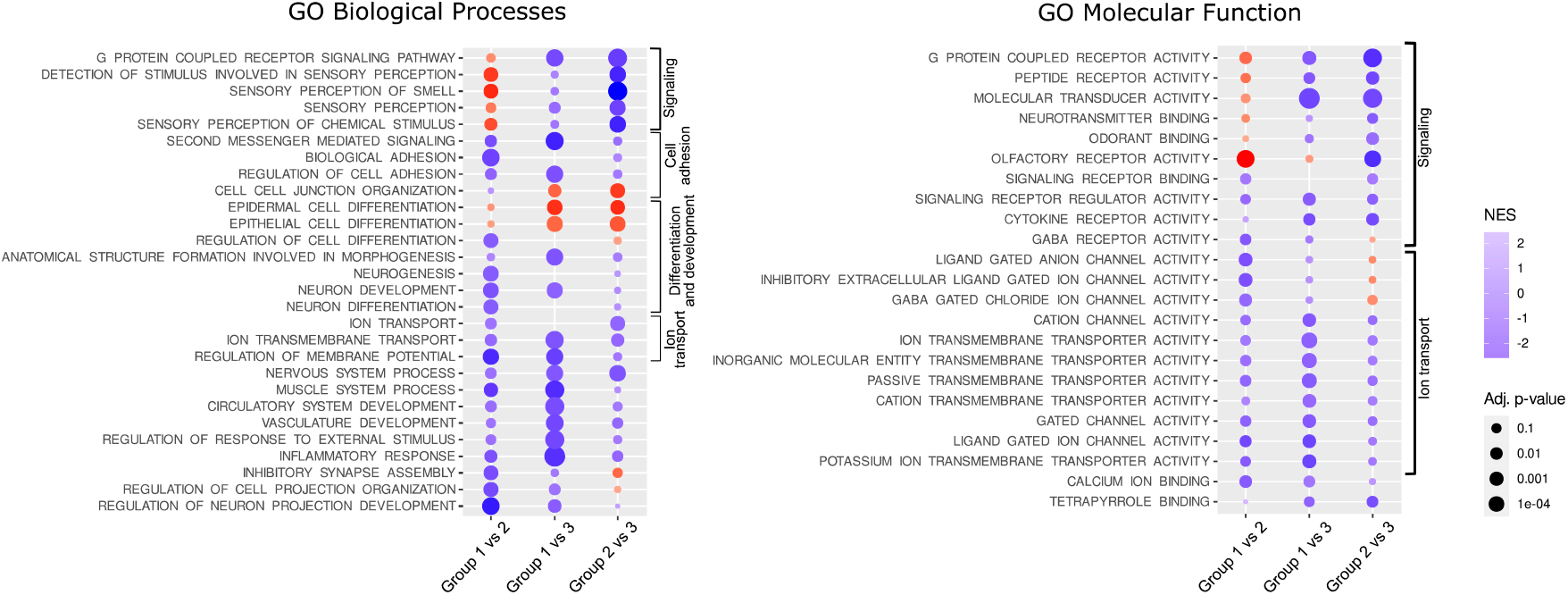
Results of Gene Set Enrichment analysis. NES – normalized enrichment score

### 3.3 Differences in the expression of genes involved in GH secretion pathway

With a given high number of DEGs we paid a special attention to those that are related to GH secretion. According to literature data growth hormone secretion is primarily induced by hypothalamic somatoliberin (GHRH) but also by ghrelin and GIP through activation of the corresponding membrane receptor on somatotroph cells. Interestingly, we found that three identified transcriptomic groups of somatotroph PitNETs differ significantly in the expression of genes coding for each receptor. Tumors within groups 1 and 2 have high expression of *GHRHR* (somatoliberin receptor), but lower expression of *GHSR* (ghrelin receptor) than tumors in group 3. Tumors in group 1, additionally to high *GHRHR* expression, have notably higher expression of *GIPR* than the two remaining transcriptomic groups. Differences were also found among genes of somatostatin and dopamine receptors that play a role in modulation of somatotroph secretory activity. Expression of *SSTR3* was at notably lower level in group 2, while a significant decrease of *SSTR5* in group 1 was observed as compared to groups 2 and 3. Higher expression of dopamine receptors *DRD1* and *DRD2* was in turn found in group 3. Irrespectively to receptor activation, Ca^2+^ influx through voltage-gated Ca^2+^ channels (VGCCs) also plays an important role in GH secretion (Nussinovitch, 2018). *CACNA1C* and *CACNA1D* encoding VGCCs were identified as expressed at significantly higher level in groups 1 and 2 as compared to group 3 (Figure 2B). The signaling pathways downstream key receptors (cAMP and Phospholipase C pathways) are mediated by many proteins that are encoded by the genes that were differentially expressed between three transcriptomic groups. They include genes encoding G-proteins: *GNAI1, GNAI2*, adenyl cyclase *ADCY1, ADCY3, ADCY4, ADCY5, ADCY7*; cAMP response elements *CREB1, CREB3L1, ATF2, ATF5B, ATF4, CREB, CREBP, EP300*; phospholipases C: *PLCB2, PLCB4, PLCB1*; protein kinases C: *PRKCA, PRKCD, PRKCE, PRKCI, PRKCZ* as well as inositol trisphosphate receptor *ITPR3*. Details are presented in Supplementary Figure 2A, while comprehensive representation of the expression levels of genes involved in GH secretion is presented in Figure 2B, which is based on KEGG pathway (visualized with pathview, original picture available in Supplementary Figure 2B). *GH1* encoding GH was also found differentially expressed, with the highest expression in transcriptomic group 1 and the lowest in group 3 (Figure 3A).

**Figure 3.**
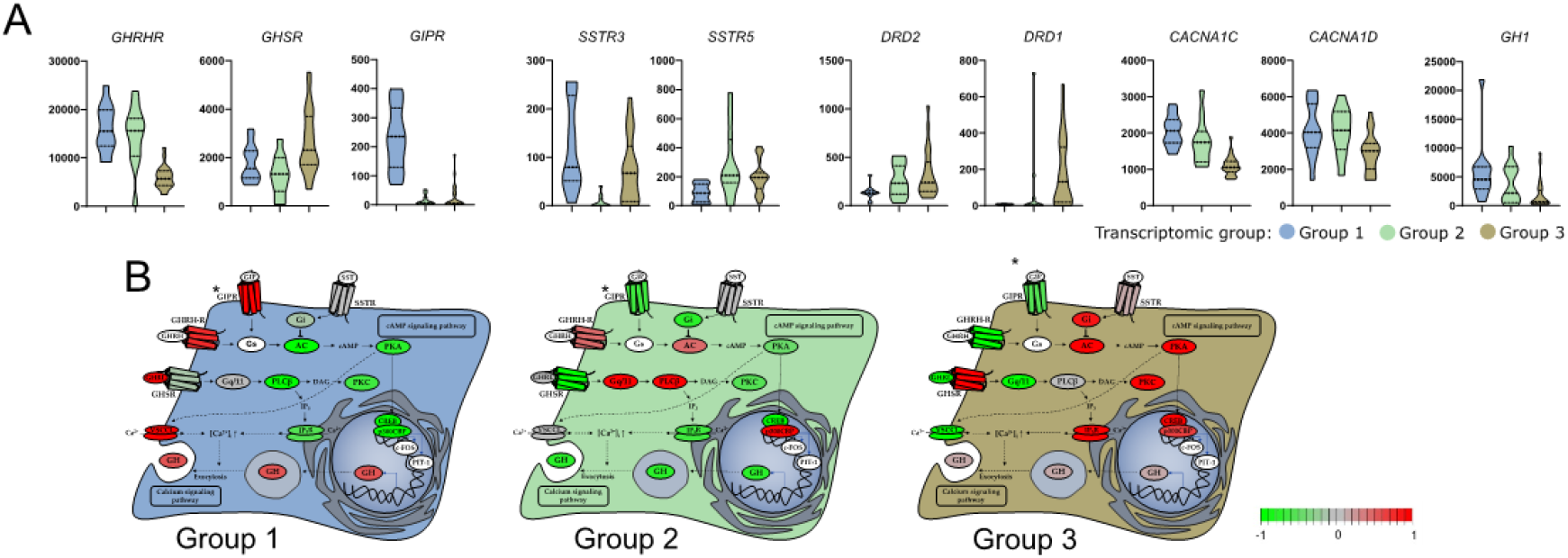
Difference in the expression of genes involved in growth hormone (GH) secretion pathway in three subtypes of somatotroph tumors. A. Expression levels of differentially expressed genes coding for cell surface receptors involved in GH synthesis and secretion. Distribution of normalized RNA-seq read counts is presented. B. Scaled normalized RNAseq read counts of genes encoding each element of GH synthesis pathway were visualized on KEGG pathway with pathview. *GIPR receptor was added manually, according to literature data (Regazzo *et al*, 2020).

### 3.4 Differences in the expression of known genes involved in somatotropinoma clinical outcome

Aberrant gene expression was previously shown to be involved in acromegaly patients’ outcome and response to SRLs. Beside the role of somatostatin receptors, the role of genes involved in epithelial-mesenchymal transition, cell proliferation and cell signaling was previously reported (Gil *et al*, 2021a) Transcriptomic groups of somatotroph PitNETs differed in the expression of genes related to epithelial-mesenchymal transition (EMT) that have proven role in acromegaly including *CDH1* (Kiseljak-Vassiliades *et al*, 2015; Chauvet *et al*, 2016), *SNAI2* (Mendes *et al*, 2018), *FLNA* (Coelho *et al*, 2019), *ARRB1* (Gatto *et al*, 2016, 2013), *RORC* (Gil *et al*, 2022) and *ESRP1* (Chauvet *et al*, 2016) but also in other genes with an important role in EMT including *CDH2, CDH3, CDH11, CTNNB1, CLDN1, CLDN3, CLDN4, CLDN9* and *ZAEB1* (Figure 4A). Differences between transcriptomic groups were also observed in expression levels of proliferation-related genes *CCND1* (Vitali *et al*, 2014), *CDKN1B* (Kiseljak-Vassiliades *et al*, 2015) and *MKI67* (Gil *et al*, 2021b) and genes involved in cell signaling *TGFB1* (Li *et al*, 2021)and *STAT3* (Zhou *et al*, 2015) that all have a reported role in acromegaly patients outcome (Figure 3B).

**Figure 4.**
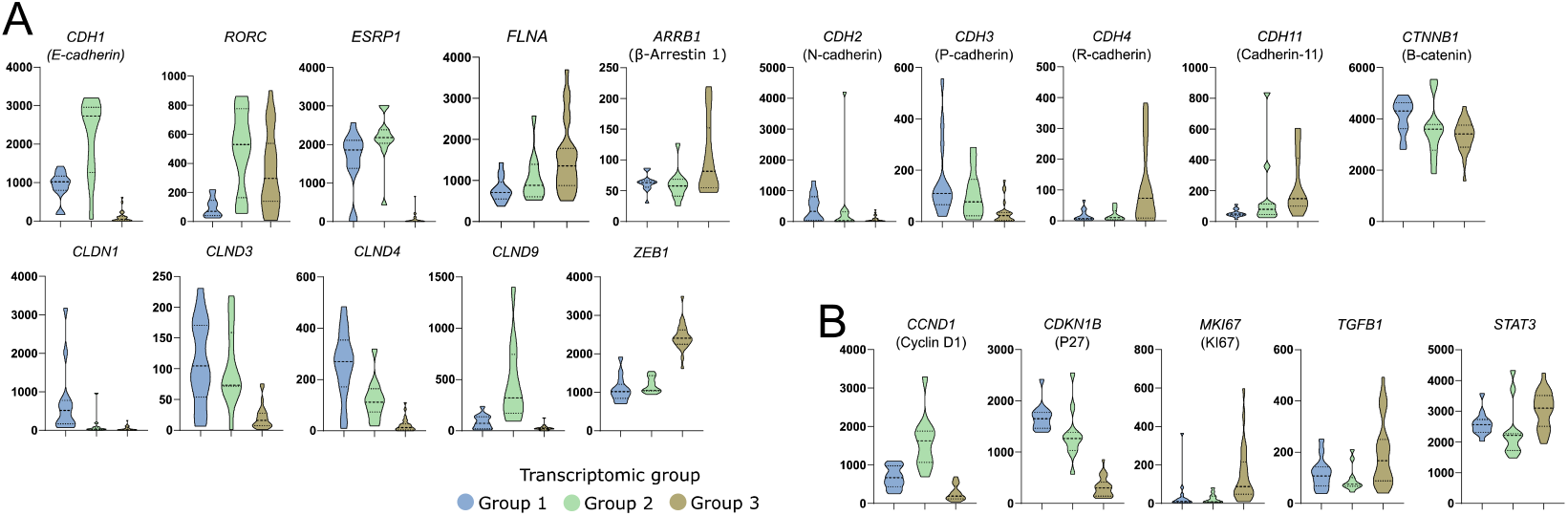
Differences in the expression of known somatotroph pituitary tumor-related genes in three subtypes of somatotroph tumors. Distribution of normalized RNA-seq read counts are presented. A. Genes encoding proteins involved in epithelial-mesenchymal transition. B. Genes related to somatotroph pituitary tumor growth.

### 3.5 Role of SF-1 (NR5A1) transcription factor in a subtype of somatotroph tumors

We explored the expression levels of known genes encoding transcription factors specific to particular lineages of anterior pituitary cells in three identified transcriptomic groups of somatotroph tumors. A striking difference in the expression level of *NR5A1* (SF-1) was observed. It is expressed on very high level in tumors from group 1 as compared to other tumors. SF-1 (*NR5A1*) is a commonly accepted marker of pituitary gonadotroph cell lineage, therefore, its higher expression in group 1 of somatotropinomas requires special attention (Figure 4). We measured *NR5A1* expression in somatotroph tumors of transcriptional group 1 and gonadotroph PitNET samples (tumor samples from our previous investigation (Rusetska *et al*, 2021)) with qRT-PCR. This showed that the range of *NR5A1* expression in group 1 somatotroph PitNETs and gonadotroph PitNETs is similar (Figure 4B). Tumors in group 1 had also significantly higher expression of *GNRHR* and *FSHB* than in other groups but no differences were found in *LHB* and *CGA* genes that are expressed in pituitary gonadotrophs. This group also has the highest level of *PRLHR* and *TRHR*, encoding other receptors of hypothalamic hormones.

Transcriptomic group 1 tumors are those with a high *GIPR* expression. GIPR was previously found to be involved in steroidogenesis process. It induces expression of known steroidogenesis-related genes including *NR5A1, STAR* and *CYP11A1* (Bates *et al*, 2012; Fujii *et al*, 2014). As expected, higher expression level of these genes was found in group 1 tumors as compared to each of the other groups. Accordingly, a significant correlation of the expression levels of *GIPR* and each of *NR5A1, STAR* and *CYP11A1* was found, with a highest correlation coefficient in *GIPR* and *NR5A1* analysis (Spearman R= 0.785, p < 0.0001) (Figure 4C).

**Figure 4.**
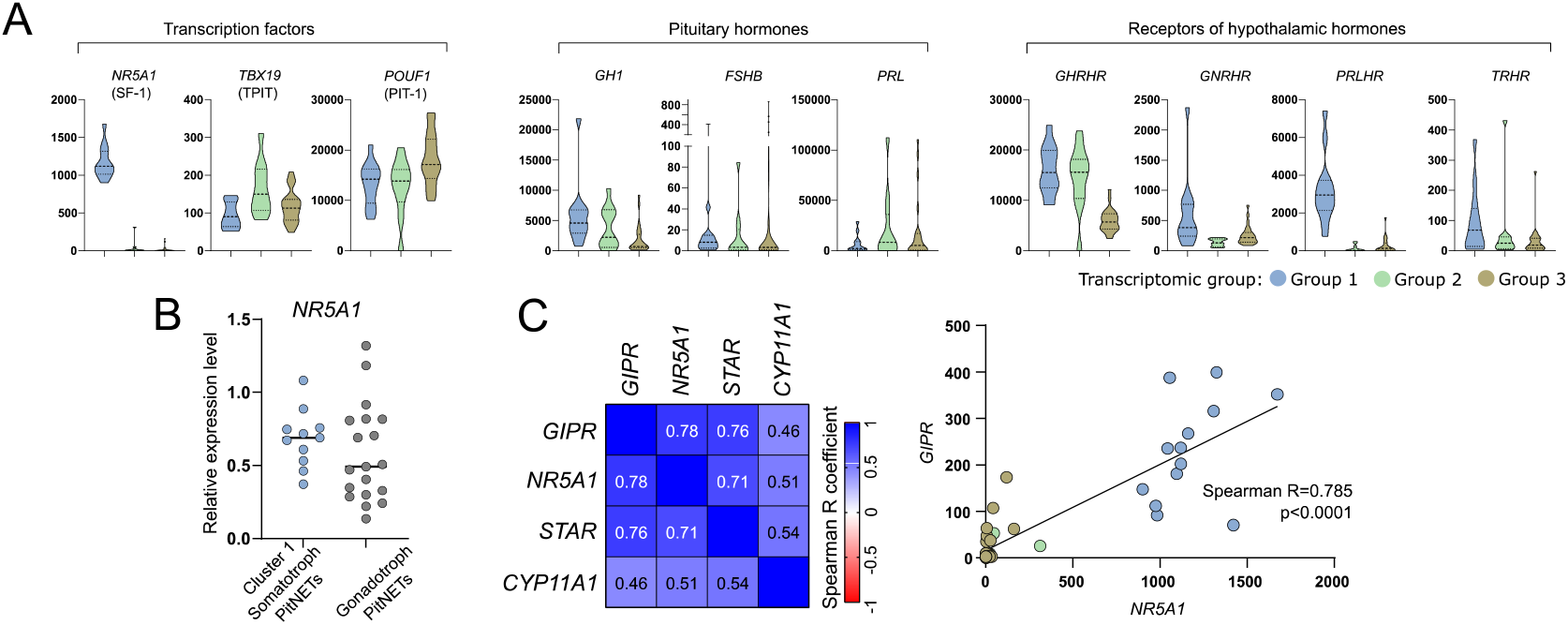
The expression of differentially expressed genes (DEGs) that are related to identity of anterior pituitary cells. A. Differential expression of transcription factors specific for anterior pituitary lineages, pituitary hormones and receptors. *POUF1* (PIT-1) was not identified as DEG based on whole transcriptome analysis, it its presented as a key transcription factor of somatotroph lineage. Difference in *POUF1* expression was determined with Kruskal-Wallis test (p=0.0074) by comparing normalized read count values; B. The comparison of *NR5A1* (SF-1) expression in somatotroph and gonadotroph PitNETs based on qRT-PCR measurement; C. Co-expression of GIPR and steroidogenesis-related genes that were found as differentially expressed in comparison of tumors form transcriptional group 1 with groups 2 and 3. Normalized RNA-seq read counts were analyzed

### 3.6 Difference in clinical/histopathological features between transcriptomic groups of somatotroph tumors

To evaluate the differences in clinical parameters between the transcriptomic groups of somatotroph tumors we used tumor samples from the entire cohort of acromegaly patients, without any intentional preselection. First, we determined whether the transcriptomic groups can be identified by the expression level of the marker genes that could be easily determined with qRT-PCR. Based on RNA-seq results we selected 9 genes that could serve as potential classifiers. Using qRT-PCR we measured the expression level of 9 potential markers in 48 samples that were previously included in transcriptome profiling and that were clearly categorized (Figure 5A). Using ROC curve analysis we determined the value of qRT-PCR-measured expression levels as classifiers of each transcriptomic group. The results of the evaluation of 9 marker genes are presented in Supplementary Figure 3. We selected three marker genes: *NR5A1, CCND2* and *SEC23A* that met criteria of area under curve (AUC) >0.99 and that allowed for selecting a clear threshold value. *NR5A1* distinguished between transcriptomic group 1 and groups 2/3, while *CCND2* and *SEC23A* discriminated groups 1/2 and group 3 (Figure 5B). The use of thresholds for *NR5A1, CCND2* and *SEC23A* expression level determined on data from RNA-seq group allows for clear categorization of nearly entire patients’ cohort (Figure 5C). Six patients (4,5%) of the entire patient cohort could not be assigned to any transcriptomic category due to a low expression level of each marker. These 6 patients were excluded from further analysis of clinical data.

**Figure 5.**
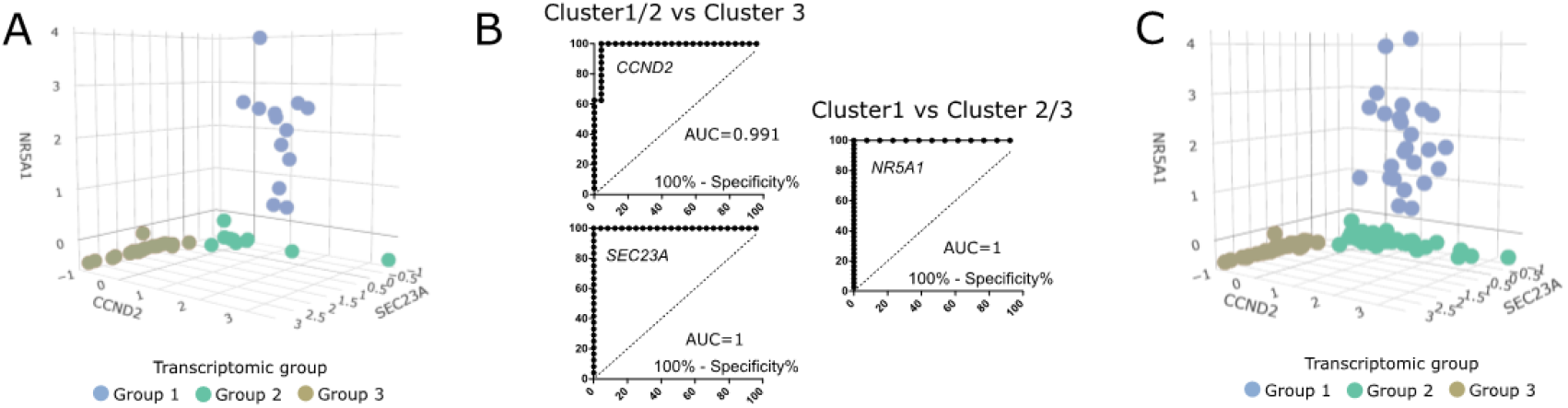
The value of classification based on three expression markers *NR5A1, CCND2* and *SEC23A*. A. The expression levels of marker genes in 48 samples categorized based on RNA-seq data. Scaled qRT-PCR expression values are presented. B. ROC curve analysis of three selected marker genes. C. The values of marker gene expression in entire patients’ cohort, with classification of the samples. 6 unclassified tumors were excluded. Scaled qRT-PCR expression values are presented.

The analysis of clinical data showed that transcriptomic group 1 which is composed of tumors positive for *NR5A1* expression are the *GNAS*-mutation negative. This group includes 25/128 patients (19,5%). Most of the tumors within this group were DG somatotroph PitNETs, however, 5 of the tumors were positive for both GH and LH upon immunohistochemical staining and were classified as plurinominal GH/LH tumors. Transcriptomic group 2 accounted for 59/128 patients (46,1%). It included vast majority of *GNAS*-mutated patients (66.1% were *GNAS*-mutated), mainly those determined as densely granulated. This group included DG somatotroph PitNETs, but also mixed GH/PRL tumors. Transcriptomic group 3 included 44/128 patients (34,4%) with low incidence of *GNAS* mutations (18% were *GNAS*mut). The difference in proportions of *GNAS*mut patients between transcriptomic groups was significant (Chi square test p<0.0001). Group 3 was composed mostly of sparsely granulated tumors as determined with electron microscopy, diagnosed mainly as SG somatotroph PitNETs. It also included 7 DG tumors (10%), but no mixed or plurinominal PitNETs. In this group, a higher proportion of invasive tumors (43%, 19/44) was observed, as compared to transcriptomic groups 1 and 2 (16.6% and 20,3%, respectively) (Chi square test p=0.0154). Data are presented in Figure 6A. Tumors from transcriptomic group 2 were significantly smaller than those from two other groups (Figure 6B). No differences were observed in serum GH and IGF-1 (Figure 6B) or patients’ demographical data (age and gender) between the groups.

**Figure 6.**
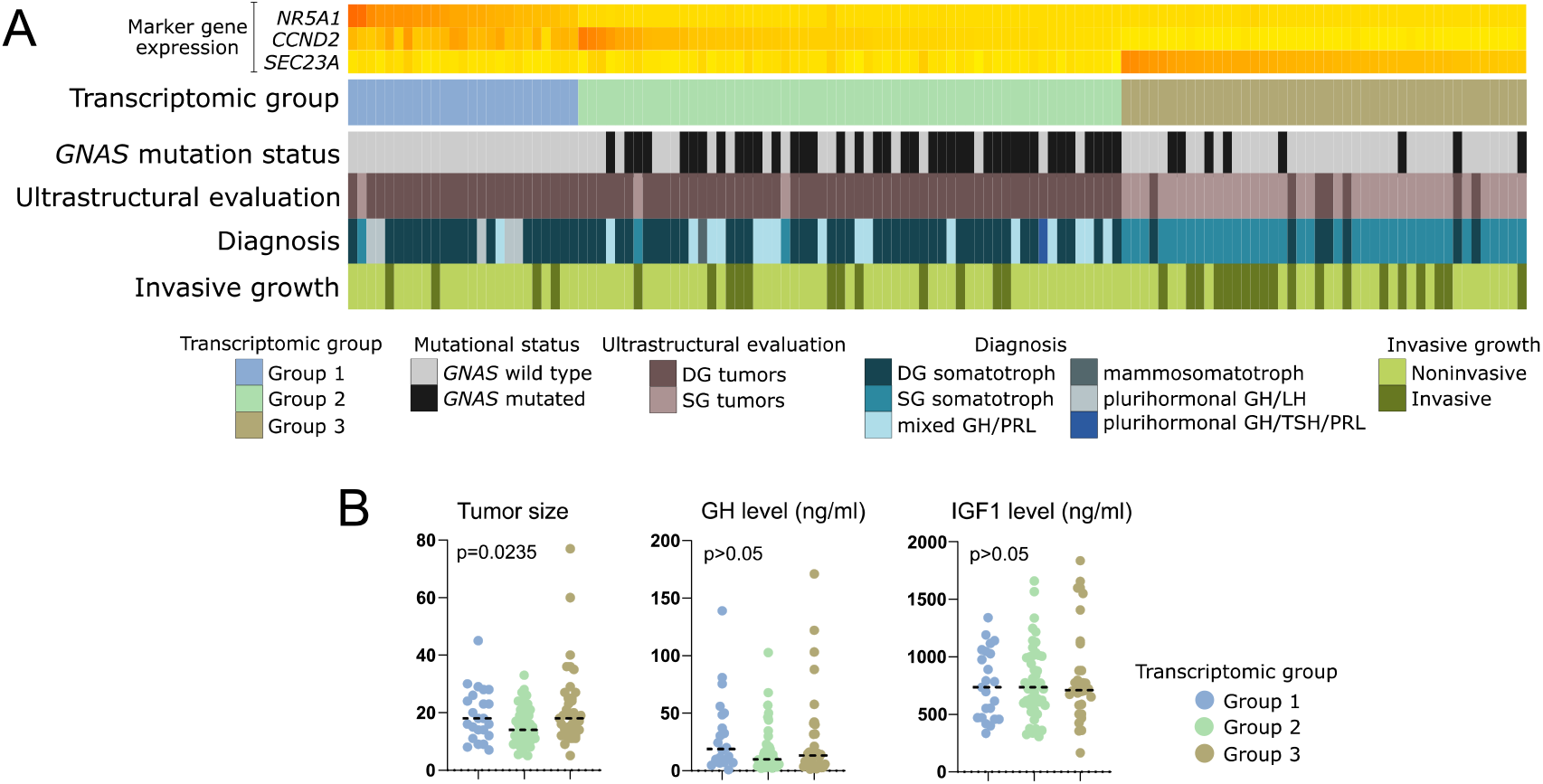
Clinical data of patients with somatotroph PitNETs categorized according to expression of marker genes into three transcriptomic groups. A. Presentation of categorical patients characteristics. B. Comparison of quantitative patients data including tumor size (maximal tumor diameter) as well as GH and IGF1 plasma level. Horizontal dashed line indicates median. P-value assessed with Kruskal-Wallis test.

**Figure 7.**
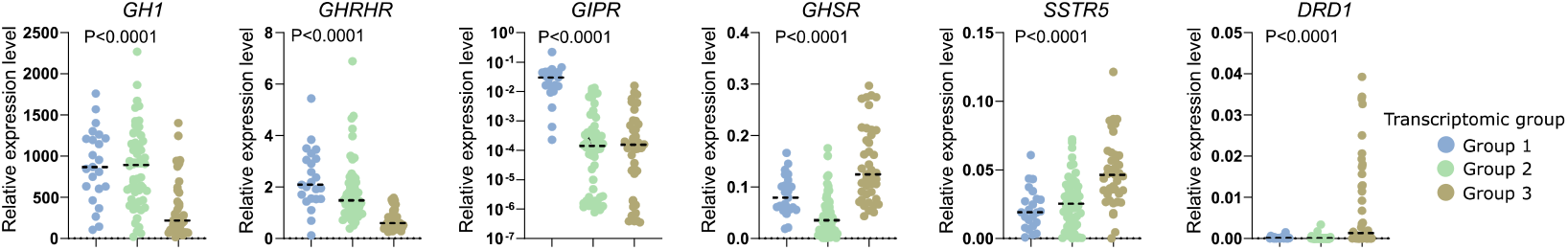
Relative expression level of key regulatory genes involved in somatotroph cell functioning measured with qRT-PCR in the entire patients‘ cohort, with patients stratified according to three transcriptomic groups. Horizontal dashed line indicates median.

RNA-seq results revealed that transcriptomic groups differ in the expression level of key genes involved in GH synthesis and secretion including *GHRHR, GIPR, GHSR, SSTR* and *DRD1* as well as *GH1* encoding GH itself. Using qRT-PCR we measured expression level of these genes in the entire patients’ cohort categorized into three transcriptomic groups according to marker genes evaluation. Significant differences in the expression of each gene were found and confirmed the observation in whole transcriptome analysis. Groups 1 and 2 have high level of *GH1* and *GHRHR*. Tumors in group 1 have high *GIPR* expression, while those in group 3 have higher expression of ghrelin receptor (*GHSR*) and *DRD1*. Slight discrepancy was observed between qRT-PCR results of the entire cohort and data from RNA-seq in terms of *SSTR5* expression level. PCR-based measurement showed the highest level of *SSTR5* expression in transcriptomic group 3.

## Discussion

The symptoms of acromegaly result from GH oversecretion by pituitary tumors which develop from PIT-1-positive anterior pituitary cell lineage. Our results show that profile of gene expression clearly discriminates somatotroph tumors into three transcriptomic groups that differ in the expression level of large number of genes. According to GSE results the expression differences are mainly related to process of cell signaling including G-protein signaling and ion transport, both of which play a key role in regulation of secretory activity of pituitary cells (Fletcher *et al*, 2018; Lania & Spada, 2009) as well as to cellular adhesion and differentiation.

Importantly, DEGs include genes directly involved in regulation of GH secretion like those coding for receptors of somatoliberin, gastric inhibitory polypeptide and ghrelin, receptors for dopamine and somatostatin as well as voltage-gated calcium channels. It seems that secretory activity in tumors of each transcriptomic subgroup may be driven by a slightly different mechanism. First group is characterized by high expression of *GHRHR, GIPR* and *VGCCs* as well as the highest expression of GH-encoding gene. The second group includes tumors which seem to be mainly related to hypothalamic GHRH stimulation, as they show high *GHRHR* expression. Tumors in the third group, in turn, are probably dependent on ghrelin signaling since they present with the highest expression of gene encoding ghrelin receptor.

*GIPR*, which is highly expressed in group 1 tumors, encodes the receptor of glucose-dependent insulinotropic polypeptide (GIP) that has a well-known role in neuroendocrine tumors (Regazzo *et al*, 2020). Upon ligand binding GIPR activates coupled heterotrimeric G-protein complex containing stimulatory Gs subunit and leads to activation cAMP pathway. Therefore, GIPR activation is considered to be mimicking GHRHR stimulation by hypothalamic GHRH hormone (Regazzo *et al*, 2020). Acromegaly patients with GIPR expression commonly react to oral glucose load with a paradoxical increase in GH level, which resembles the induction of hormone secretion in food-dependent Cushing's syndrome (Hage *et al*, 2021). DNA methylation profiling in *GIPR*-high *versus GIPR*-low somatotroph PitNETs showed a difference in genome-wide methylation pattern which is in line with our results indicating that this group represents a separate molecular subtype (Hage *et al*, 2019). Interestingly, high expression of *NR5A1* encoding SF-1 transcription factor was found exclusively within this transcriptional group. According to classification of PitNETs, SF-1 is a well-established marker of gonadotroph tumors (Asa *et al*, 2022b), and the observation that it is also expressed in a particular subtype of somatotroph tumors may suggest a need of slight revision of classification criteria. We observed that *GIPR*-high somatotroph PitNETs express *NR5A1* at the level comparable to gonadotroph tumors. The expression of SF-1 in some somatotroph PitNETs was also recently noticed by Mario Neou et al. (Neou *et al*, 2020). In our study we clearly demonstrate that somatotroph tumors with SF-1 expression are exactly those that express *GIPR*. It appears to be functionally related. *GIPR* plays a role in steroidogenesis, as previously observed in adrenal cortex-derived cell line and mice. The experiments showed that manipulating *GIPR* expression results in changes of transcription levels of *NR5A1, STAR* and *CYP11A1* (Bates *et al*, 2012; Fujii *et al*, 2014). Transcriptomic group 1 tumors are those with significantly highest expression of *GIPR, NR5A1, STAR* and *CYP11A1* as compared to tumors from other groups. Accordingly, correlation analysis indicates that these genes are co-expressed with *GIPR*. This suggests that a high expression level of *NR5A1* in transcriptomic group1 of somatotroph tumors results from high GIPR-related signaling. The mechanism underlying high *GIPR* expression in somatotroph tumors was the matter of previous research and it is still unclear (Hage *et al*, 2019).

Beside the expression of SF-1, tumors in transcriptomic group 1 have higher expression of *GHRHR* (gonadoliberin receptor) and *LHB* (LH) than other transcriptomic subtypes which may suggest that they may share some molecular features with gonadotroph tumors. It should be noted however that no difference in the expression of *FSHB* or *CGA* (subunit α) was observed between groups, and group 1 has also the highest level of receptors for other hypothalamic hormones (PRLHR and TRHR).

We are convinced that transcriptomic classification of acromegaly-causing tumors may have clinical implications. The use of SRLs is basic pharmacological treatment for acromegaly patients. Therapy response and prognosis were previously found to be related to expression of somatostatin receptor genes (*SSTR2* and *SSTR5*) (Kiseljak-Vassiliades *et al*, 2015; Iacovazzo *et al*, 2016), genes involved in EMT (Gil *et al*, 2021a), and those involved in cell proliferation like *CDKN1B* (p27) (Kiseljak-Vassiliades *et al*, 2015)or *MKI67* (Ki-67) (Gil *et al*, 2021b). Loss of cadherin E was clearly linked with a poor response to SRLs (Puig-Domingo *et al*, 2020). Importantly, striking differences in the expression of *CDH1* were observed between transcriptomic groups with the highest expression in group 2 and the lowest in group 3. The expression profile of other genes, that were previously shown to be related to SRLs response,(Kiseljak-Vassiliades *et al*, 2015; Chauvet *et al*, 2016; Gatto *et al*, 2016; Gil *et al*, 2022; Iacovazzo *et al*, 2016) suggests also that tumors in transcriptomic group 2 may be the most sensitive to somatostatin analogs. They have a favorable pattern of gene expression - those related to EMT (high *RORC* and *ESRP1* and low *ARRB1* (β-arrestin)) as well as *SSTR5* (the highest expression among subtypes), *MKI67* (low expression) and *CDKN1B* (high level).

Preliminary analysis of clinical data from samples included in RNA-seq showed that transcriptomic profile corresponds to pathomorphological diagnosis. In general, transcriptomic group 1 includes *GNAS*wt, densely granulated somatotroph PitNETs, group 2 includes *GNAS*wt and *GNAS*mut DG tumors, and transcriptomic group 3 includes predominantly SG tumors. Unfortunately, the proportions between the pathomorphological subtypes of somatotroph tumors among the samples included in our RNA-seq do not reflect those in general population of acromegaly patients. Therefore, we made an attempt to evaluate the true proportions in patients’ numbers in individual groups and clinical significance of transcriptomic classification in a large group of acromegaly patients, without any intentional preselection. We evaluated candidate marker genes that allow for stratification of patients into three transcriptomic subtypes and used three most potent markers to classify the entire patients’ cohort. This classification is based on 3 markers was not adequate only in 6 patients that were subsequently excluded from the analysis. The results allow to draw some general conclusions. They clearly show that in transcriptomic group 1 there are patients with *GNAS*wt, DG tumors (mainly pure somatroph PitNETs but also tumors that are positive for both GH and LH) with low frequency of invasive growth. These tumors are positive for both *NR5A1* (SF-1) and *GIPR* expression and probably include the patients with paradoxical GH response to glucose intake, according to previous studies (Hage *et al*, 2021). These patients are the least frequent and they account for approximately 20% of patients suffering from acromegaly. Transcriptomic group 2 seems to be the most numerous (46% of acromegaly patients). It includes mostly *GNAS*mut, densely granulated tumors as determined with electron microscopy, diagnosed generally as DG somatotroph PitNETs, and additionally mixed GH/PRL tumors. The smaller tumor size found in this group is concordant with the observation that a favorable gene expression profile was observed in these tumors. Accordingly, low rate of invasive growth was reported for these patients. This corresponds to general observation of better prognosis in patients with densely than sparsely granulated tumors (Kontogeorgos *et al*, 2022). In the third transcriptomic group there are sparsely granulated tumors with low frequency of *GNAS* mutations diagnosed mainly as SG somatotroph PitNETs. They account for approximately 35% of tumors causing acromegaly. These tumors have the highest rate of invasive growth and unfavorable gene expression profile (low expression of *CDH1, RORC* and *ESRP1* and high *ARRB1, MKI67, ZEB1, STAT3* and *TGFB1* levels). They have also high level of *DRD1* gene expression, encoding stimulatory dopamine receptor that activates cAMP pathway (Rosas-Cruz *et al*, 2021). This may affect the results of treating these patients with dopamine analogs like cabergoline. In general, the identified gene expression patterns correspond to literature data indicating that patients with sparsely granulated tumors are considered as high risk tumors (Kontogeorgos *et al*, 2022) as they have have worse prognosis, lower rate of postoperative remission, tumors with frequent invasive growth, a tendency to regrow after surgery and lower response to SRLs (Kontogeorgos *et al*, 2022). Our transcriptomic data indicate that these tumors may be significantly driven by ghrelin signaling. They have higher expression of ghrelin receptor gene as compared to densely granulated tumors. The role of ghrelin signaling in GH secretion and pathogenesis of somatotroph PitNETs was already determined (Devesa, 2021). Importantly, ghrelin receptor was recognized as a therapeutic target (Liang *et al*, 2020). Small inhibitors against this receptor are available and were already subjected to clinical trials on treatment of various human diseases (Liang *et al*, 2020). The use of this therapeutical approach may be potentially beneficial in sparsely granulated tumors with high *GHSR* expression and it could complement standard treatment with SRLs.

## Data availability

RNA-Seq data is available at ArrayExpress: E-MTAB-11889 (https://www.ebi.ac.uk/arrayexpress/experiments/E-MTAB-11889)

## Acknowledgements

This study was supported by Maria Sklodowska-Curie National Research Institute of Oncology, grant SN/GW04/2020.

The study was approved by the local Ethics Committee of Maria Sklodowska-Curie Institute - Oncology Center in Warsaw, Poland. Informed Consent was obtained from all subjects, and experiments conformed to the principles set out in the WMA Declaration of Helsinki and the Department of Health and Human Services Belmont Report.

## Author Contributions

Julia Rymuza: Conceptualization, Methodology, Formal analysis, Investigation, Visualization, Writing—review and editing; Paulina Kober: Formal analysis, Investigation, Writing—original draft preparation; Natalia Rusetska: Investigation; Beata J. Mossakowska: Investigation; Maria Maksymowicz: Resources, Data curation, Writing—review and editing; Aleksandra Nyc: Resources, Investigation; Szymon Baluszek: Writing—review and editing, Formal analysis, Visualization; Grzegorz Zieliński: Resources, Data curation; Jacek Kunicki: Resources, Data curation; Mateusz Bujko: Conceptualization, Methodology, Formal analysis, Visualization, Writing—original draft preparation, Funding acquisition

